# Neuronal innervation of breast cancer promotes metastatic dissemination

**DOI:** 10.1101/2025.11.25.690523

**Authors:** Anthony J. Schulte, Eran R. Andrechek

**Author notes:** Corresponding author Eran R. Andrechek Professor Department of Physiology Michigan State University 2194 BPS Building 567 Wilson Road East Lansing, MI 48824 (517)884-5020 Lab (517)355-5125. This work is supported by T32GM142521.

## Abstract

Metastasis is a leading cause of mortality in breast cancer patients, yet the signaling promoting metastatic dissemination is not completely understood. Prior literature implicates neuronal innervation in tumor progression, including recent studies in breast cancer progression with the 4T1 and PyMT cell line orthotopic injection models. Our experiments address the immune limitations of these studies with an alternative model to elucidate neuronal control of metastatic breast cancer by using a MMTV-PyMT transplant model and resiniferatoxin (RTX) for denervation. To this end, we generated a robust array of spontaneous MMTV-PyMT tumors with various histological subtypes. These tumors were transplanted into RTX challenged MMTV-Cre mice. In contrast to previous literature, denervation did not impact survival or tumor growth. Interestingly, we noticed a slight reduction in the percent of the solid poorly differentiated tumors with a corresponding increase in tumors that contained a mixed pathology in RTX challenged mice. Strikingly, and consistent with prior work, we noted a reduction in metastasis with denervation. Together, these data suggest neuronal innervation promotes metastasis without impacting tumor growth.

## Introduction

Metastasis is the leading cause of mortality in breast cancer patients (1). To combat this, 80% of breast cancer patients receive adjuvant therapy to preemptively eradicate possible metastases (2). However, despite this therapeutic approach, mortality occurs in 40% of patients with metastases (2). This therapeutic approach may also subject patients to unnecessary toxic side effects, ranging from cardiotoxicity (3,4) to peripheral neuropathy (5,6). It is therefore critical to understand the driving factors of breast cancer progression and metastasis to develop more precise therapies to increase survival and reduce toxic side effects in patients with breast cancer metastasis.

To study breast cancer metastasis, there are a number of model systems, but one of the most commonly employed is the MTMV-PyMT genetically engineered mouse. This mouse line rapidly develops multifocal mammary tumors with an average onset of 45 days due to expression of the Polyoma Middle T antigen in the mammary epithelium. Importantly, these mice develop pulmonary metastasis in the majority of the mice (7).

Neuronal innervation is crucial in mammary gland development (8) as well as regulating mammary stem cell activity (9). In addition to normal development, neuronal innervation has been implicated in the metastatic progression of breast cancer (10). To study innervation, capsaicin has frequently been used as a potent agonist of TRPV1 receptor, resulting in the ablation of neurons (11–13). TRPV1 is a calcium channel expressed primarily on sensory neurons, along with some immune cells. Continuous administration of capsaicin has been shown to reduce metastasis and tumor burden in the syngeneic 4T1 model (11). Similarly, chemical-induced neuronal ablation has increased survival and decreased progression of pancreatic ductal adenocarcinoma (12). Given that the 4T1 study was limited to a single model and that there were immune concerns, we sought to determine if neuronal control of metastatic breast cancer was critical in other models where the immune system was not perturbed. With PyMT being widely used to study metastasis, we sought to examine the role of neuronal innervation in metastasis. To this end, we generated MMTV-PyMT tumors and used them for viable transplantation into mice treated with resiniferatoxin (RTX). MMTV-Cre transgenic mice (14) were used as a recipient since a portion of the MMTV promoter / enhancer is transcribed and translated and is immunogenic (15). RTX is a super agonist of sensory neuron-expressed calcium channel TRPV1 (8, 9) and administration causes calcium-mediated neuronal death (6, 8). This approach allowed us to test the hypothesis that denervation of the tumor microenvironment specifically, without denervation of the tumor itself, would decrease tumor growth and prevent metastasis.

Recently, an elegant study used the TRPV1 knockout mouse model in conjunction with PyMT cell mammary orthotopic injections to test this same hypothesis (18). This, like our study, investigated the role of neuronal innervation of the tumor microenvironment in tumor growth and metastasis. However, due to this publication, we have not undertaken a more detailed analysis since our findings complemented the work from the Oudain laboratory.

## Materials and Methods

### Mice

All animal husbandry and use were in compliance with local, national and institutional guidelines. Ethical approval for the study was approved by Michigan State University Animal Care & Use Committee (IACUC) under AUF 06/18-084-00. MMTV-PyMT634 mice were obtained from The Jackson Laboratory. Mice were monitored twice weekly for tumor initiation and growth. At a 1.5 cm endpoint, mice were necropsied. Tumors and lungs were stained with hematoxylin and eosin for histological subtyping and presence of pulmonary metastases. The number of metastases was quantified using a single cut through the lung and count of the number of micro-metastases in that plane.

### TRPV1+ neuron chemo-ablation

Resiniferatoxin (RTX) was obtained from Alomone Labs (Cat # R-400). As previously described to ablate TRPV1+ neurons (13), RTX was injected subcutaneously with escalating doses (30 μg/kg at day 1, 70 μg/kg at day 2 and 100 μg/kg at day 3) into the abdominal mammary fat pad of 8 week old MMTV-Cre mice (14). Control mice received a vehicle solution of DMSO with Tween80 in phosphate-buffered saline (PBS).

### Mammary fat pad transplantation

MMTV-PyMT mammary tumors were harvested and stored in DMEM with 20% FBS and 10% DMSO at −80 °C before long term storage in liquid nitrogen vapor phase. Tumors were thawed and orthotopically implanted into the abdominal mammary gland of 12-week-old MMTV-Cre female mice. Mice were palpated twice weekly for tumor growth. When the tumor size reached 1.5 cm, mice were necropsied for further analysis.

### Histology

Samples for histology were fixed in 10% formalin and submitted to Michigan State University Pathology lab.

### Statistical analysis

Statistical significance was determined by student’s T-test or Fisher’s exact test *p<0.05

## Results

Denervation has been shown to reduce tumor burden and metastasis in the syngeneic 4T1 breast cancer model (11) as well as other cancers including PDAC (12). The MMTV-PyMT mouse model is highly metastatic (7) but it was unknown whether neuronal innervation impacts tumor growth and metastasis in this context. We hypothesized that RTX challenge would reduce tumor growth and metastasis in MMTV-PyMT tumors transplanted into MMTV-Cre mice. We generated MMTV-PyMT females and collected tumors from 3 separate MMTV-PyMT mice. The histological patterns of these tumors was diverse as previously reported (19,20), including solid, comedoadenocarcinoma, microacinar, papillary, and EMT tumors (Fig. 2A). The overall percentage of each histology is presented in Figure 2B. Of the variety of histological subtypes, tumors with mixed pathology were the predominant phenotype, with papillary also being common. These MMTV-PyMT tumors were then viably frozen for tumor transplantations.

**Figure 1.**
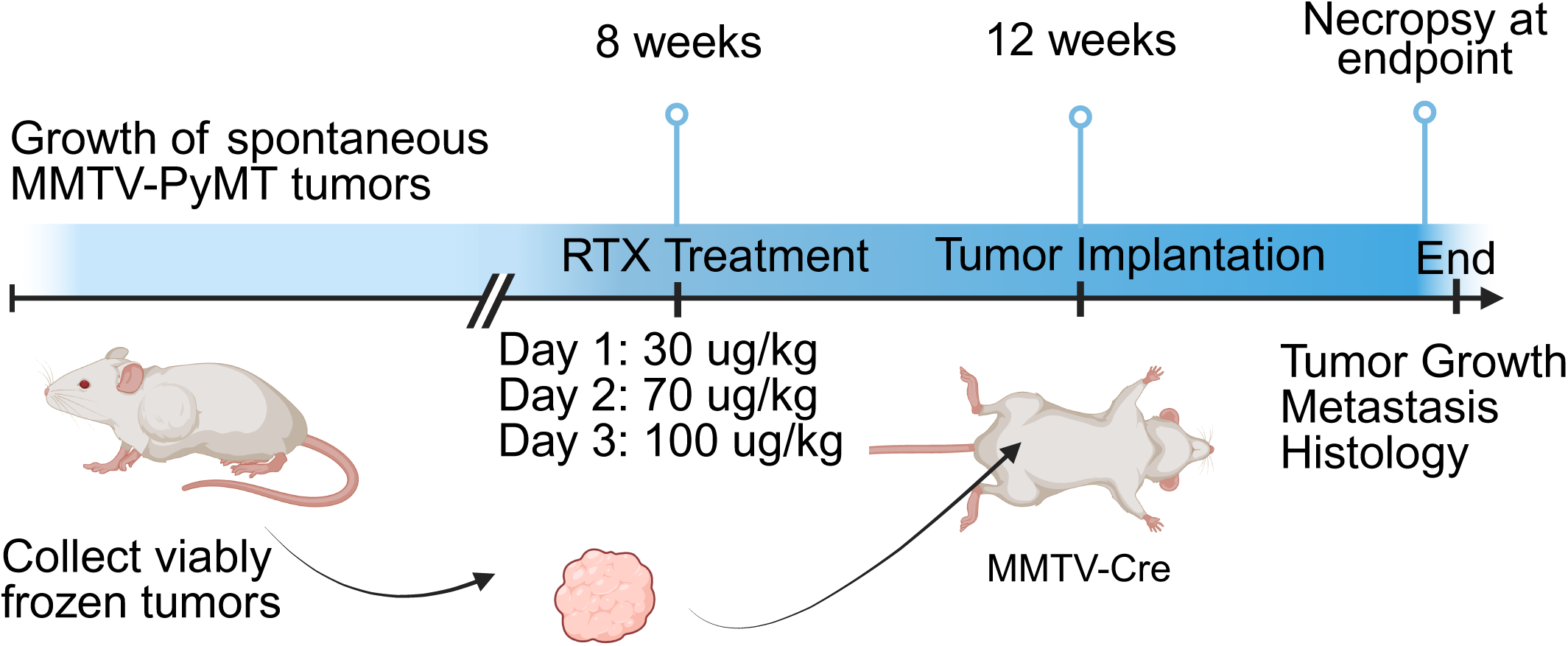
Experimental Overview. The experimental overview depicts that tumors from 3 separate MMTV-PyMT mice were collected for viable transplantation. At 8 weeks of age, MMTV-Cre mice were treated with either resiniferatoxin (RTX) or vehicle at escalating doses of 30, 70, or 100 ug/kg for 3 days. At 12 weeks, MMTV-PyMT tumors were transplanted into the mammary fat pad of the MMTV-Cre mice. Tumor growth was recorded and mice were necropsied at endpoint. Tumors were collected for histology and lungs were assessed for metastasis. Created in BioRender. Schulte, A. (2025) https://BioRender.com/oa7dz3k.

**Figure 2.**
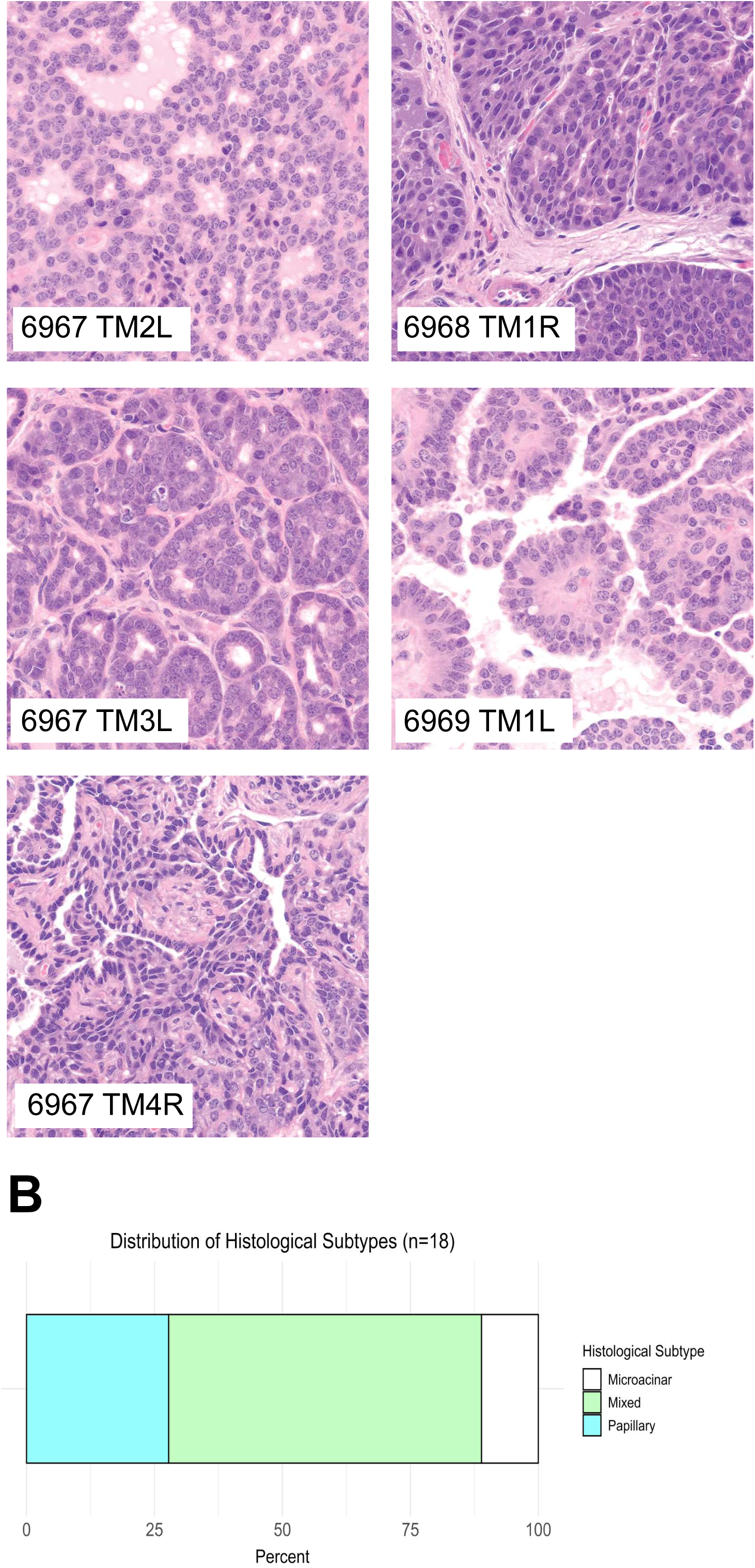
MMTV-PyMT original tumors generated are histologically diverse. H&E staining of tumors from MMTV-PyMT mice showed diverse histology, with many displaying mixed histopathology. 6967 TM2L had solid and comedoadenocarcinoma patterns. 6967 TM3L had microacinar and EMT patterns with intervening stromal tissue. 6967 TM4R was largely microacinar. 6968 TM1R and 6969 TM1L was papillary in nature (A). The percentages of histological subtypes observed from the three MMTV-PyMT mice showed a diverse array, with mixed pathologies being the most prominent at 61%, followed by papillary at 27% and microacinar at 11% out of the 18 tumors (B).

TRPV1+ sensory neurons were ablated at 8 weeks of age in MMTV-Cre recipient mice and MMTV-PyMT tumors were transplanted into these mice 4 weeks later. Survival was similar between the two groups, as shown by the Kaplan Meier plot in Figure 3A. Likewise, tumor growth was monitored twice weekly and tumor area was recorded, plotted against days (Fig. 3B). The time for individual tumors to reach 1.5cm^2^ was not significantly different (Figure 3C). The tumor growth of the vehicle and RTX treated groups was also not significantly different, suggesting that RTX treatment did not change tumor growth or survival. To assess if RTX treatment would impact the histological subtype that develops, we implanted the same original tumors into both vehicle and RTX treated mice and observed the pathologies that resulted. While no new histological subtypes were formed, we noted a slight reduction in the percental of the solid poorly differentiated tumors with a corresponding increase in tumors that contained a mixed pathology (Figure 4).

**Figure 3.**
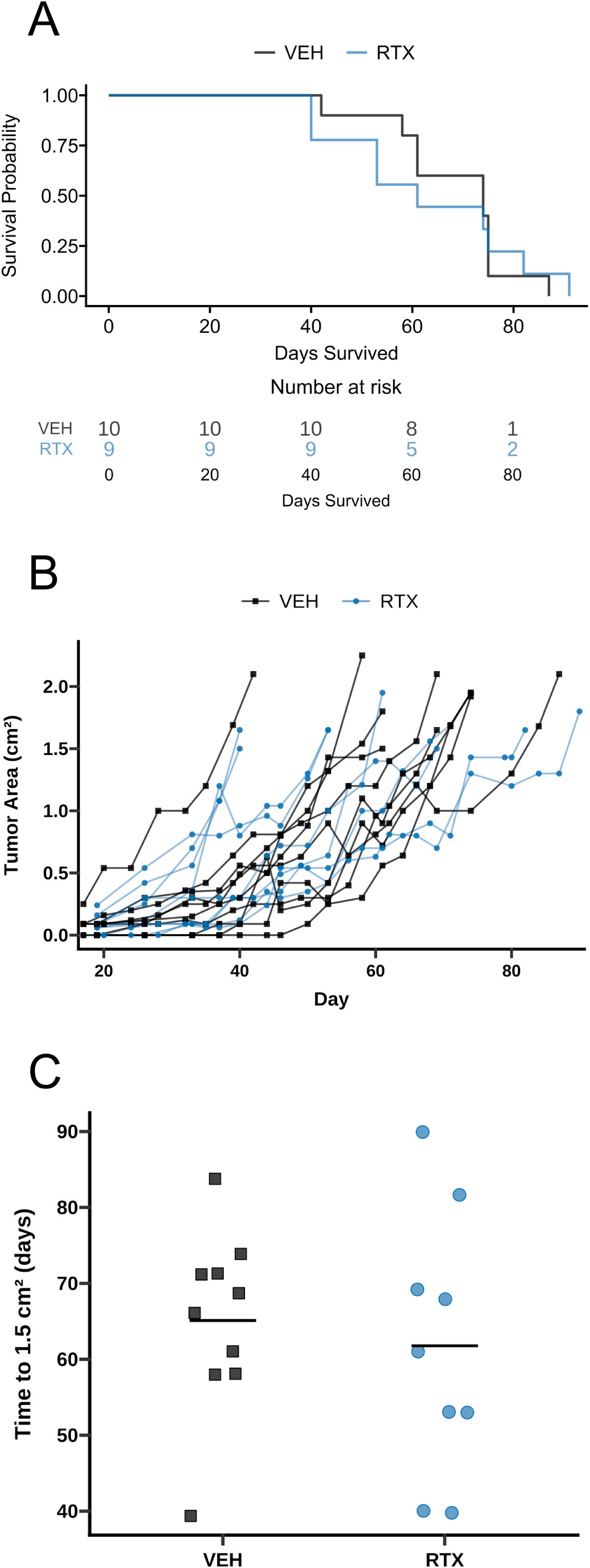
Denervation does not impact tumor growth or survival. Following tumor transplantation in RTX or VEH treated mice, tumor growth was monitored twice weekly until endpoint. Time to reach endpoint for each mouse was recorded. The probability of survival is plotted versus time in days in Kaplan-Meier curves (A). Tumor growth is plotted as tumor area vs time in days for each individual mouse (B). Time in days for tumors to reach 1.5 cm^2^ is summarized in (C).

**Figure 4.**
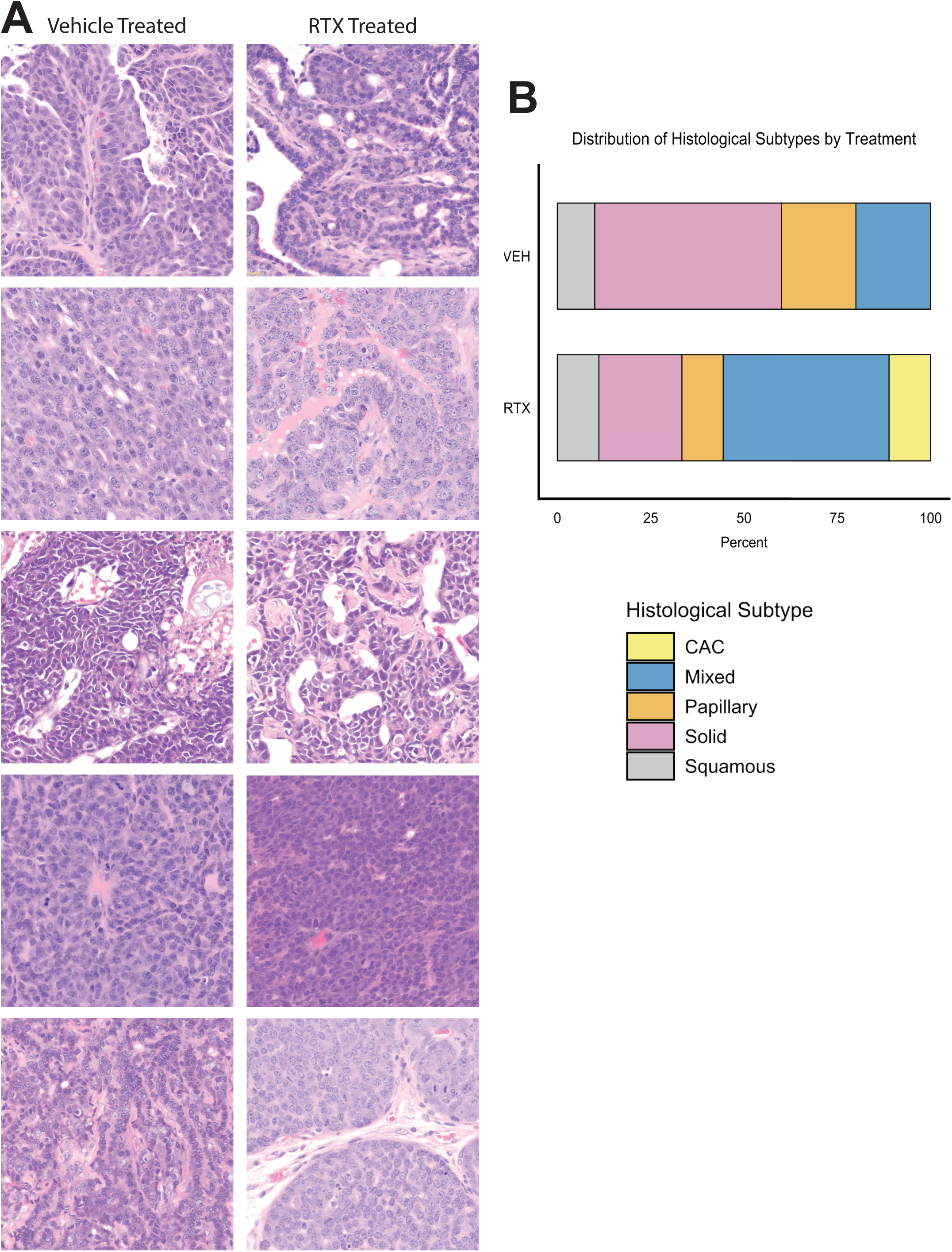
Denervation shifts histological subtype proportions. The same viable frozen tumors were implanted into vehicle and RTX treated recipient mice. The histological subtypes of the resulting tumors are shown. Both vehicle and RTX mice exhibited mixed, papillary, solid, and squamous histology (A). The histological subtypes observed are summarized by group with alterations to the subtype prevalence noted (B).

Previous studies suggested that neuronal innervation was associated with metastasis (11), (18). In support of this, when assessing pulmonary metastasis, denervated mice were noted to completely lack metastatic lesions. We examined the lungs of mice challenged with vehicle, where pulmonary metastases had formed and were readily observed (Figure 5A). In contrast, no metastases were observed in RTX challenged mice (Fig. 5B). Indeed, 4 of 10 vehicle treated mice had clear metastases while 0 of 9 mice challenged with RTX had observable metastases (Figure 5C). While a Fisher’s two-tailed t-test (p=0.087) showed this to be nearing significance, the study was somewhat underpowered. However, the absence of metastasis in conjunction with the literature suggests denervation reduces metastasis in this MMTV-PyMT transplant model.

**Figure 5.**
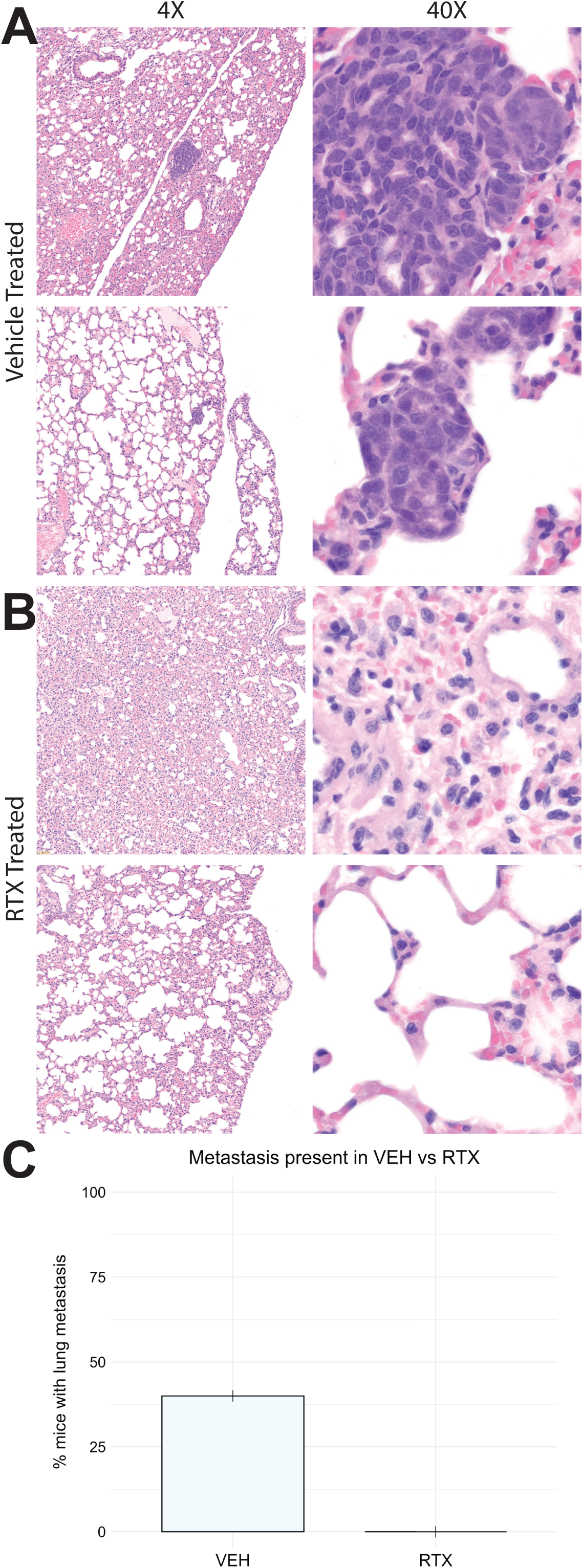
Neuronal innervation permits metastasis in MMTV-PyMT tumor transplants. Metastasis was observed in H&E staining of lungs in VEH treated mice with matched 4X and 40X samples demonstrating the metastases (A). In contrast, no metastasis was observed in RTX treated mice (B). Quantification revealed a metastasis rated of 40% in control vehicle treated mice (n=10) and no metastasis in RTX treated mice (n=9). Results were analyzed with a two-tailed Fisher’s exact test, p=0.087 (C).

## Discussion

The innervation of mammary tumors has been hypothesized to regulate tumor growth and metastatic spread through a number of potential mechanisms. To study the role of innervation in this process in mice with an intact immune system, we ablated sensory neurons and then implanted metastatic tumors into the mammary fat pad. This resulted in changes to primary tumor histology and reduced metastatic progression using the PyMT model system. This work is complementary to recent findings using the PyMT model in a cell line approach (18).

Given that both human and mouse mammary glands, as well as tumors, are known to be highly innervated (11), elegant studies have shown that denervation via capsaicin or genetic knockout decreased tumor growth and metastasis in the 4T1 syngeneic model (11) and recently in a PyMT cell line (PY8119) model system (18) respectively. Numerous studies in other tissues have also implicated neuronal innervation in tumor growth and metastasis. Indeed, it is known that pancreatic and prostate tumors use innervation to grow and metastasize (12,21). However, one limitation to many of these studies that ablate TRPV1 is that this protein is also expressed on several immune cell types (22–24). Thus, the observation of anti-tumor effects may be confounded by reduced pro-tumor immunity. To address this, we used RTX administration to permanently kill sensory neurons prior to tumor transplant. A 4-week recovery period allowed recovery of immune cells, abolishing this confounding variable. Furthermore, by denervating prior to tumor transplantation, we were able to test the role of sensory neurons specifically in the tumor microenvironment.

Previous studies used cell lines to investigate the role of neuronal innervation in tumor growth and metastasis. However, breast cancer is a heterogenous disease and using a cell line does not allow this diversity to be captured. By employing a MMTV-PyMT transplant model, the tumors with various gene expression profiles (20) allowed us to examine the role of innervation on tumor histology. Indeed, various histological subtypes were generated from MMTV-PyMT mice and while tumors were implanted into both vehicle and RTX treated recipients, we noted a shift in tumor histopathology. Interestingly, despite the histological alterations, tumor growth and time to endpoint were similar between mice challenged with RTX and vehicle (Fig. 3). This is contradictory to prior literature, in which denervation decreases tumor burden (11) or tumor weight (18). This deviance could be due to cytotoxic immune effects, tumor intrinsic denervation, and altered genomic profiles from different original tumors. Overall, our results suggest that denervation of the tumor microenvironment is insufficient to reduce tumor growth and improve survival in the MMTV-PyMT transplant model.

The primary goal of this study was to examine the role of innervation in mediating metastatic progression. We noted that metastasis was reduced in RTX challenged mice relative to vehicle controls. This recapitulated denervation studies in both 4T1(11) and MMTV-PyMT (18) models. Interestingly in the 4T1 model, capsaicin was continuously administered to produce cytotoxic effects to sensory neurons. In the PY8119 model, a TRPV1 global knockout model was used. However, as TRPV1 is expressed in T-cells (22), macrophages (23), and eosinophils (24), this presents additional variables, complicating the differentiation of effects of denervation on tumor growth and metastasis from effects on immune cytotoxicity. Indeed, these methods may confound effects of tumor-intrinsic neuronal signaling relative to signaling from the tumor immune microenvironment. By using RTX to ablate sensory neurons prior to tumor implantation, we were able to discern the effects of denervating only the microenvironment and demonstrate that metastasis was indeed impacted, although this was not quite statistically significant. Further investigation into what signaling is altered through denervation of the tumor immune microenvironment with this model should be considered to uncover new signaling driving metastasis. The interplay between the two is especially intriguing given that other studies have examined requirements of the immune system in metastatic progression in this model. For instance, macrophage recruitment has been shown to be crucial for metastatic progression in MMTV-PyMT mice (24,25). With the RTX / PyMT model, as immune cells have time to recover and repopulate following RTX treatment, the study of neuro-immune interaction that drive tumor progression may be of interest. Moreover, by using 5 different spontaneous tumors derived from 3 different MMTV-PyMT mice, this robustness permits us to conclude this is not a clonal effect.

Taken together, our results support the recent study from the Oudain laboratory demonstrating that PyMT mouse mammary tumors require innervation for metastasis. Use of an RTX model and immunocompetent recipient mice have demonstrated that metastasis reduction is an innervation specific effect and the alteration of the tumor histopathology suggests that signaling pathways within the primary tumor have also been altered.

## Notes

### Competing Interest Statement

The authors have declared no competing interest.

### Funding

- nih: R01CA160514, T32GM142521

